# High-Throughput, automated assay for detection of colonization by *Candida auris*

**DOI:** 10.64898/2026.09.01.748466

**Authors:** Francesca Scala, Brandon Schumitsch-Jewell, Diane Podzorski, Hannah Gander, Megan Lasure, Alana K. Sterkel

## Abstract

*Candida auris* is an emerging multidrug-resistant fungal pathogen associated with healthcare-associated outbreaks, persistent colonization, and invasive infections. Increasing demand for surveillance has created a need for high-throughput methods capable of supporting large-scale screening programs. We developed and validated an automated laboratory-developed real-time PCR assay for detection of *C. auris* colonization on the Hologic Panther Fusion® open-access platform and compared its performance with the existing BD MAX™ assay.

Analytical performance was evaluated by assessing limit of detection, accuracy, precision, specificity, inclusivity, reproducibility, and reagent and specimen stability. The Panther Fusion® assay demonstrated a limit of detection of approximately 18 CFU/reaction and showed 97% overall agreement with the BD MAX™ assay. Positive and negative percent agreement were 94% and 100%, respectively, with excellent agreement between methods (κ = 0.94). No cross-reactivity was observed with non-*C. auris* organisms, all five major *C. auris* clades were detected, and assay performance remained stable across operators, reagent and specimen storage conditions.

Following implementation, 26,838 clinical specimens were tested on the Panther Fusion® platform. Retrospective analysis demonstrated lower equivocal (0.28%) and indeterminate (0.09%) rates than those observed on the ABI and BD MAX™ platforms. Among PCR-positive specimens that underwent culture, the Panther Fusion® assay demonstrated 87.24% culture agreement. Because retrospective data were collected during different testing periods and patient populations, comparisons between platforms were not used to assess relative assay sensitivity or specificity. Implementation of the Panther Fusion® assay increased surveillance testing capacity from approximately 88 to 500 specimens per shift while maintaining robust analytical performance.

**IMPORTANCE:** *Candida auris* continues to pose a significant public health challenge, as evidenced by the growing demand for surveillance testing. This study demonstrates that the Panther Fusion® assay provides a high-throughput, automated approach for *C. auris* surveillance that substantially increases testing capacity while maintaining accurate and reproducible detection. By enabling large-scale screening while maintaining optimal analytical performance, this assay can help laboratories meet increasing surveillance demands and support timely public health and infection prevention efforts.

---

*Candida auris* is an emerging multidrug-resistant fungal pathogen that has become a major public health concern because of its ability to cause healthcare-associated outbreaks, persistently colonize patients, and cause invasive infections associated with substantial morbidity and mortality (1–3). Environmental persistence and efficient transmission within healthcare settings distinguish *C. auris* from many other *Candida* species and contribute to its ability to establish prolonged outbreaks in hospitals and post-acute care facilities (1,4). Colonized individuals serve as important reservoirs for transmission, making rapid identification of asymptomatic carriers a critical component of infection prevention and outbreak control programs (5,6).

In the United States, both clinical and screening detections of *C. auris* have increased substantially over the past decade, prompting expansion of surveillance activities by healthcare facilities and public health agencies (2,5). Recent national surveillance data identified more than 21,000 individuals with positive colonization screening results during 2016–2023, underscoring the growing demand for large-scale screening programs and laboratory capacity to support them. Screening programs are particularly important in long-term acute-care hospitals, skilled nursing facilities, and other high-risk healthcare settings where ongoing transmission can occur despite infection control interventions (1,5,6).

Culture-based screening methods remain important for isolate recovery and downstream characterization; however, molecular assays provide substantially faster turnaround times and improved scalability for surveillance efforts. Leach et al. developed a highly sensitive real-time PCR assay capable of detecting *C. auris* directly from surveillance specimens and demonstrated its utility for rapid public health response. Subsequent investigations during the New York outbreak confirmed the value of PCR-based screening for large-scale surveillance and outbreak management. More recently, several clinical and public health laboratories have reported successful implementation of laboratory-developed PCR assays for *C. auris* colonization screening, highlighting the continued need for adaptable, high-throughput molecular testing approaches (6–8).

To support national surveillance efforts, the CDC Antimicrobial Resistance Laboratory Network established regional reference laboratories that provide specialized testing for *C. auris*. As the Midwestern regional reference center, the Wisconsin State Laboratory of Hygiene (WSLH) performs colonization screening for multiple states and processes large volumes of surveillance specimens (Figure 1). The WSLH previously performed *C. auris* testing using the BD MAX™ platform, which provided automated testing but limited overall throughput. To increase testing capacity and reduce hands-on labor, we developed and validated a laboratory-developed real-time PCR assay on the Hologic Panther Fusion® open-access platform. A recent study demonstrated successful implementation of a Fusion-based *C. auris* assay within a healthcare system outbreak response program, but data describing implementation within a high-volume regional public health reference laboratory remain limited. Accordingly, we evaluated the analytical performance, reproducibility, stability, and implementation characteristics of a Panther Fusion® *C. auris* assay designed to support large-scale public health surveillance testing. (7).

**FIG 1.**
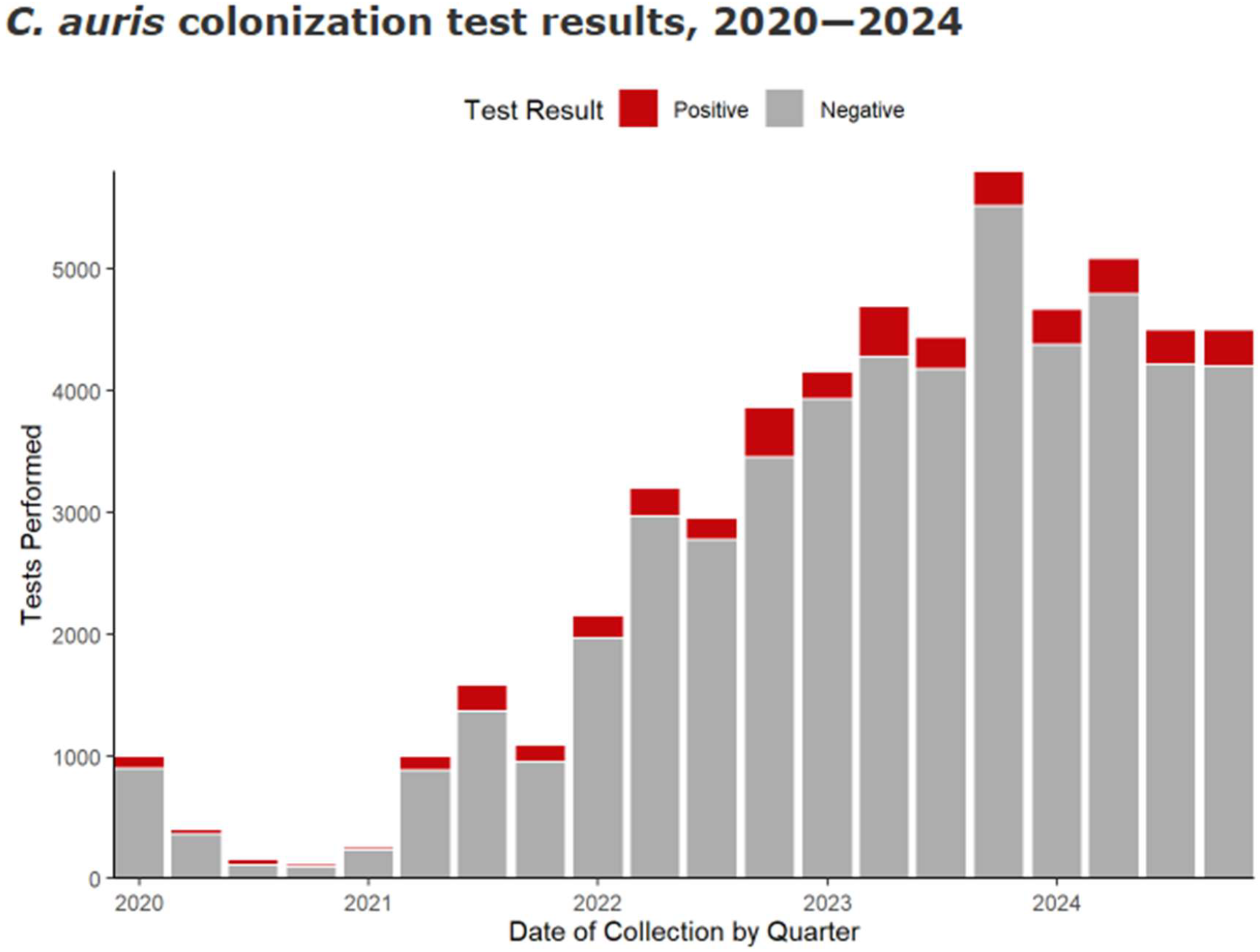
Annual Candida auris surveillance testing volume at the Wisconsin State Laboratory of Hygiene from 2020–2025. Increasing testing demand prompted implementation of higher-throughput molecular testing platforms.

## MATERIALS AND METHODS

Validation studies were performed in accordance with Clinical Laboratory Improvement Amendments (CLIA) requirements and College of American Pathologists (CAP) guidelines for laboratory-developed molecular assays. Analytical studies included assessment of sensitivity, specificity, inclusivity, reproducibility, precision, and stability prior to implementation for clinical testing.

### Clinical specimens

Residual surveillance swab specimens submitted for *Candida auris* colonization screening were used for assay validation and implementation studies. Specimens were collected using standard surveillance collection procedures and transported in liquid Amies medium. Residual deidentified specimens remaining after routine clinical testing were used under institutional policies governing quality improvement and laboratory assay validation activities.

For method comparison studies, 100 residual surveillance specimens previously tested by the BD MAX™ *C. auris* assay were selected, including 50 positive and 50 negative specimens. Specimens were stored at 2 to 8°C until testing on the Panther Fusion® platform.

### Culture methods

Specimens were cultured according to Wisconsin State Laboratory of Hygiene standard operating procedures (9). Aliquots were inoculated onto selective fungal media and incubated under appropriate conditions for recovery of *C. auris*. Suspect colonies were identified using matrix-assisted laser desorption ionization–time of flight mass spectrometry (MALDI-TOF MS). Culture results were used as a comparator for selected validation and retrospective analyses.

### Panther Fusion® *Candida auris* Assay

The assay was developed on the Hologic Panther Fusion® Open Access platform (Hologic, Marlborough, MA). Primer and probe sequences targeting the internal transcribed spacer 2 (ITS2) region of the ribosomal operon were adapted from the assay described by Leach et al. (2018). The assay was configured as a laboratory-developed test on the Panther Fusion® Open Access platform using Panther Fusion® Extraction Reagents-X. Default thermocycler parameters were used, and the Ct threshold was lowered to 500. Each reaction included an internal amplification control supplied by the manufacturer. Samples with quantification cycle (Cq) values <40 for the *C. auris* target were interpreted as positive. Specimens with no target amplification and acceptable internal control amplification were interpreted as negative. Indeterminate and equivocal interpretations were assigned according to predefined assay criteria.

### Analytical sensitivity

Analytical sensitivity was assessed using serial 10-fold dilutions of a cultured *C. auris* isolate prepared from a quantified stock suspension. Dilutions were tested in parallel on the Panther Fusion® and BD MAX™ platforms. Multiple replicates were evaluated at each concentration level. The limit of detection (LOD) was defined as the lowest concentration consistently detected across replicates.

### Accuracy and method comparison

Accuracy was evaluated using 100 residual clinical specimens previously tested by the BD MAX™ assay. Positive percent agreement (PPA), negative percent agreement (NPA), positive predictive value (PPV), negative predictive value (NPV), and overall agreement were calculated. Cohen’s kappa coefficient was used to assess agreement between methods.

For specimens with available culture results, Panther Fusion® results were compared with culture findings.

### Precision and reproducibility

Precision was evaluated by repeated testing of a positive specimen over 15 nonconsecutive days. Mean Cq values standard deviations (SD), and coefficients of variation were calculated.

Inter-operator reproducibility was assessed by three independent operators testing identical specimens on three separate days. Qualitative agreement and Cq SD variation were evaluated.

### Inclusivity and analytical specificity

Analytical inclusivity was assessed using representative isolates from all five major *C. auris* phylogenetic clades: Clade I AR 0382, Clade II AR 0381, Clade III AR 0383, Clade IV AR 0385, Clade V AR 1097. Detection of each isolate was confirmed on the Panther Fusion® platform.

Analytical specificity was evaluated using a panel of potentially cross-reactive organisms, including closely related *Candida* species and microorganisms commonly encountered in surveillance specimens. Isolates utilized for analytical specificity analysis included *Staphylococcus aureus, Acinetobacter baumanii*, Candida *duobushaemulonii* (AR-0391) *Candida duobushaemulonii* (AR-0392) *Candida haemulonii* (AR-0395) *Candida haemulonii* (AR-0393) *Candida tropicalis* (AR-0345) *Candida duobushaemulonii Candida glabrata* (AR-0314) *and Candida albicans* (AR-0762). Cross-reactivity was defined as generation of a falsepositive *C. auris* result.

### PPR Mix and specimen stability

The Primer and Probe Reagent (PPR) mix stability was assessed following storage under multiple conditions, including refrigerated storage, frozen storage, and extended onboard instrument storage. Performance was evaluated using positive and negative control materials.

Specimen stability was assessed by storing selected specimens in Panther Fusion® specimen lysis buffer and retesting after predefined storage intervals. Specimens were well mixed after thawing. Cq values obtained after storage were compared with baseline results.

### Retrospective implementation analysis

To evaluate assay performance during routine clinical implementation, retrospective testing data generated between 2020 and 2025 were extracted from the laboratory information system. Data from three PCR platforms were analyzed: Applied Biosystems® 7500 Fast Dx (ABI), BD MAX™, and Panther Fusion®.

Result frequencies were calculated for positive, negative, equivocal, and indeterminate categories. For PCR-positive specimens that underwent culture, culture agreement was determined by calculating the proportion of culture-positive specimens among all cultured PCR-positive specimens.

Because culture was not routinely performed on PCR-negative specimens and because testing platforms were used during different time periods and surveillance populations, retrospective data were not used to estimate clinical sensitivity, specificity, PPV, or NPV.

### Statistical analysis

Descriptive statistics were calculated using Microsoft Excel. Agreement statistics included PPA, NPA, PPV, NPV, overall agreement, and Cohen’s kappa coefficient. Exact or Wilson 95% confidence intervals were calculated where appropriate. Continuous variables are reported as means ± standard deviations.

## RESULTS

### Assay development and analytical sensitivity

The *Candida auris* assay was successfully implemented on the Panther Fusion® Open Access platform using primers and probe targeting the ITS2 region. Analytical sensitivity was evaluated using serial 10-fold dilutions of a quantified *C. auris* isolate. All dilution levels were detected by the Panther Fusion® assay, with mean Cq values increasing as organism concentration decreased (Figure 2). The lowest concentration tested corresponded to approximately 18 CFU per reaction. At this concentration, all Panther Fusion® replicates were detected, while one BD MAX™ replicate failed detection, establishing an analytical limit of detection of approximately 18 CFU/reaction.

**FIG 2.**
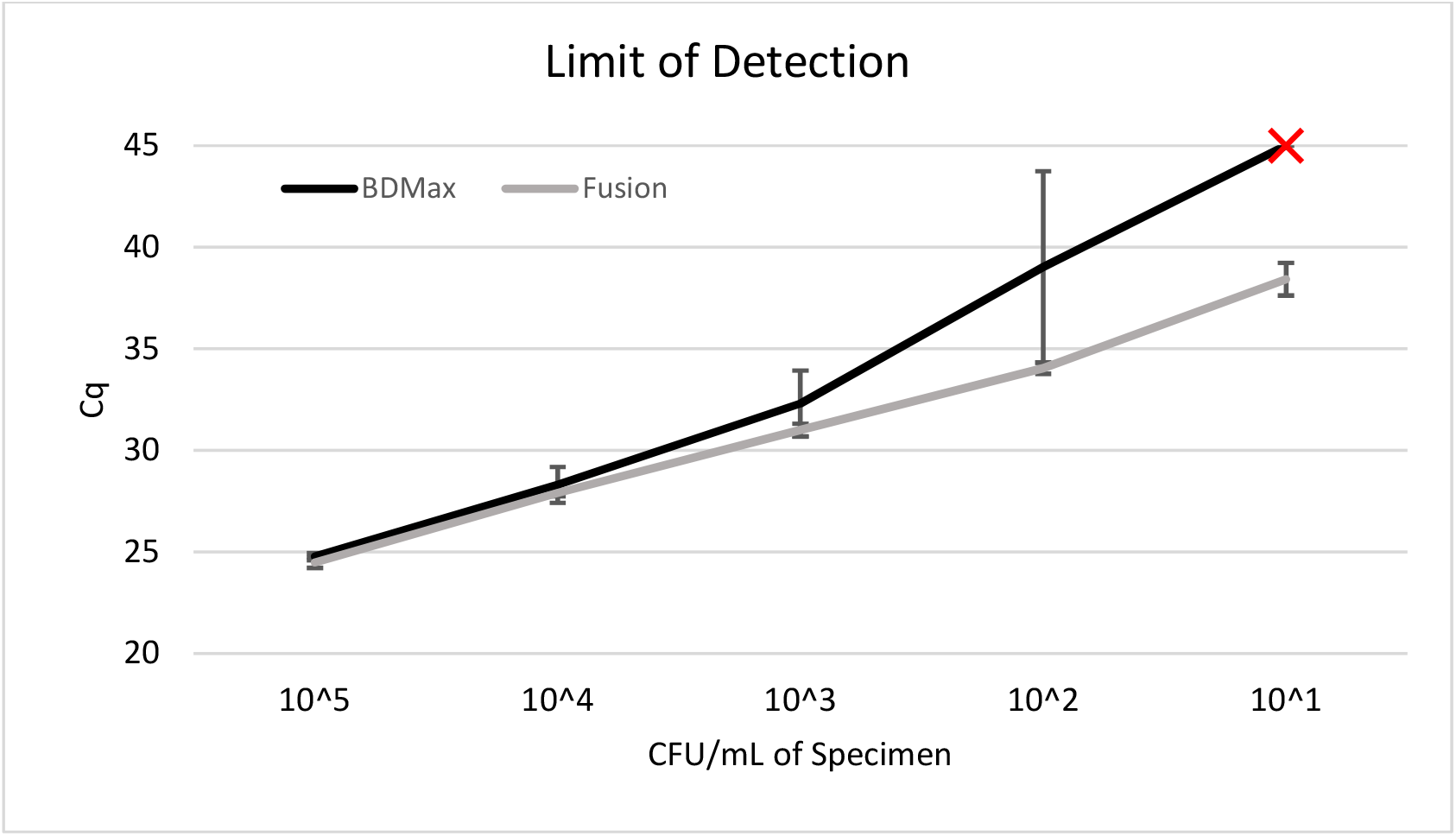
Detection of serial dilutions of C. auris by Panther Fusion® and BD MAX™ assays. Error bars represent standard deviations of replicate testing.

### Accuracy and method comparison

Accuracy was evaluated using 100 residual surveillance specimens previously tested by the BD MAX™ assay, consisting of 50 positive and 50 negative specimens. Panther Fusion® correctly identified 47 of 50 BD MAX™-positive specimens and all 50 BD MAX™-negative specimens (Figure 3). The Fusion assay detected 48 of 50 culture positive specimens and agreed with all the culture negative specimens.

**FIG 3.**
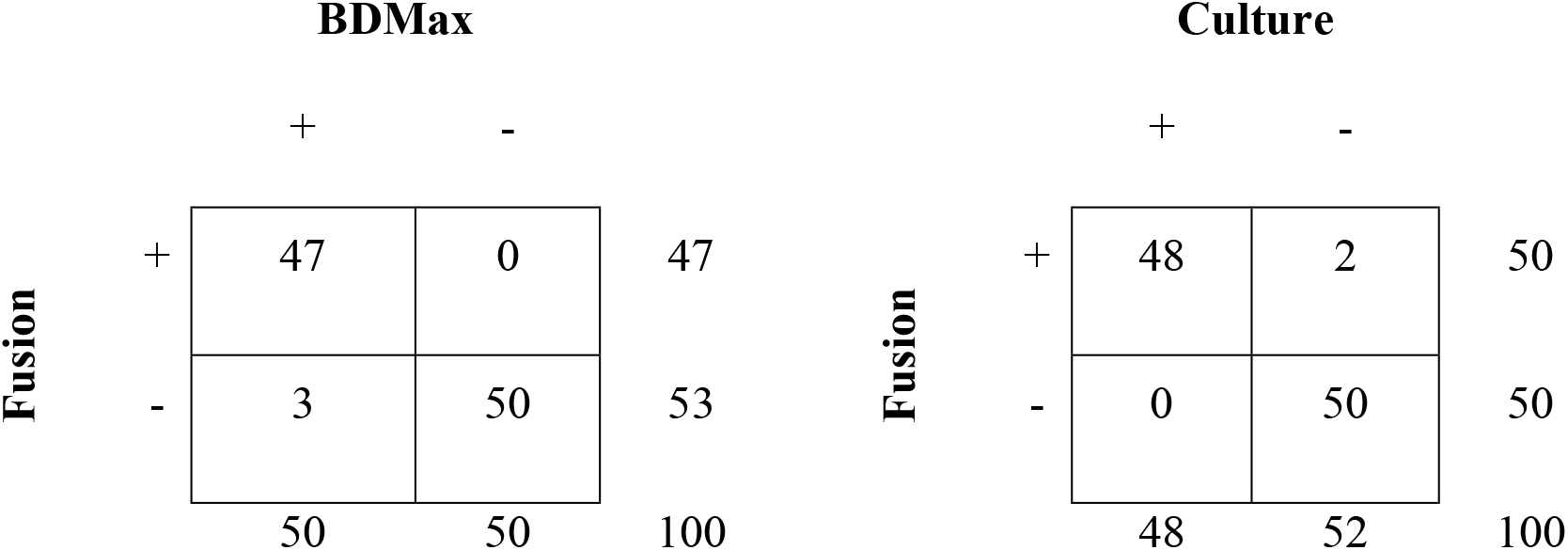
Summary of Fusion Assay performance compared to BDMax assay and culture.

Overall agreement between the two PCR methods was 97.0% (97/100). Positive percent agreement was 94.0% (47/50; 95% CI, 83.8%–97.9%), and negative percent agreement was 100.0% (50/50; 95% CI, 92.9%–100.0%). Positive and negative predictive values were 100.0% (95% CI, 92.4%–100.0%) and 94.3% (95% CI, 84.6%–98.1%), respectively. Agreement between assays was excellent (Cohen’s κ = 0.94). Comparison with culture demonstrated detection of all culture-positive specimens by Panther Fusion®. Positive and negative percent agreement with culture were 100% and 96%, respectively (Table 1).

**TABLE 1.** Summary of assay performance. PPA and NPA compared to BD Max.

| Parameter | Result |
| --- | --- |
| LOD | 18 CFU/reaction |
| PPA | 94% |
| NPA | 100% |
| Kappa | 0.94 |
| Inclusivity | 5/5 clades |

### Precision and reproducibility

Assay precision was assessed by repeated testing of a positive specimen over 15 days. The mean Cq value was 32.64 with a standard deviation of 0.45 cycles, corresponding to a coefficient of variation of 1.4%. All replicates produced positive results.

Reproducibility was evaluated by three operators testing identical specimens on three separate days. All expected positive and negative results were correctly identified by each operator, resulting in 100% qualitative agreement (15/15 concordant results). For positive specimens, Cq values showed minimal variability between operators, with standard deviations ranging from 0.31 to 0.64 cycles.

### Inclusivity and analytical specificity

The assay detected representatives of all five major *C. auris* phylogenetic clades. No false-negative results were observed during inclusivity testing.

Analytical specificity was evaluated using a panel of non-auris Candida species and other microorganisms potentially present in surveillance specimens. No cross-reactivity was observed, and all non-target organisms produced negative results.

### PPR Mix and specimen stability

PPR mix stability onboard the Panther Fusion® instrument was evaluated over 15 days (days 1, 7, 14, and 15). All time points met acceptance criteria and yielded consistent amplification results, with mean Cq values of 32.62 and 32.58 and standard deviations of 0.47 and 0.35 cycles, respectively. PPR mix refrigerated at 2 to 8°C for 30 days was evaluated on day 1, week 1, week 2, week 3, and month 1. Mean target Cq values were 32.14 and 32.20, with standard deviations of 0.34 and 0.61 cycles, respectively. Internal control amplification remained stable, with mean IC values of 28.62 cycles and standard deviations of 0.50 to 0.52 cycles. PPR mix stored at −20°C was evaluated on day 1, week 1, week 2, week 3, month 1, month 2, month 6, and month 23. Low variability was observed for both target amplification (mean Cq, 32.9 to 33.1; SD, 0.5 to 0.6 cycles) and internal control amplification (mean IC, 28.6 to 28.75; SD, 0.7 cycles), demonstrating stable performance during long-term frozen storage. Specimens stored in Panther Fusion® specimen lysis buffer at 4°C remained stable throughout the 14-day evaluation period. The mean difference between baseline and repeat-testing Cq values was 0.8 ± 1.27 cycles, and no qualitative result changes were observed.

Overall, PPR mix performance remained acceptable under all storage conditions evaluated. All target and internal control measurements were tested in duplicate.

### Retrospective implementation analysis

Between 2020 and 2025, a total of 69,226 *C. auris* surveillance specimens were tested using three molecular platforms: ABI 7500 Fast Dx (n = 26,919), BD MAX™ (n = 15,469), and Panther Fusion® (n = 26,838).

Positive results were reported in 7.53%, 8.45%, and 5.69% of specimens tested on the ABI, BD MAX™, and Panther Fusion® platforms, respectively (Table 2). Negative results accounted for 91.33%, 90.43%, and 93.95% of results, respectively. The Panther Fusion® assay generated the lowest proportion of equivocal results (0.28%) compared with ABI (1.13%) and BD MAX™ (0.82%). Indeterminate results were also less frequent on Panther Fusion® (0.09%) than on BD MAX™ (0.30%).

**TABLE 2.** Distribution of positive, equivocal, and indeterminate C. auris PCR results by testing platform. Rates were calculated from routine surveillance specimens tested between 2020 and 2025.

|  | ABI | BDMAX™ | Fusion® |
| --- | --- | --- | --- |
| Tests | 26,929 | 12,469 | 26,838 |
| Positive | 7.53% | 8.45% | 5.69% |
| Equivocal | 1.13% | 0.82% | 0.28% |
| Indeterminate | 0.01% | 0.30% | 0.09% |

Among PCR-positive specimens that underwent culture, culture-positive recovery was observed in 1,547 of 2,026 ABI-positive specimens (76.4%), 890 of 1,307 BD MAX™-positive specimens (68.1%), and 1,258 of 1,442 Panther Fusion®-positive specimens (87.2%) (Table 3). Culture positivity among equivocal specimens was uncommon across all platforms, occurring in 9.21% (28/304), 9.45% (12/127), and 6.67% (5/75) of equivocal ABI, BD MAX™, and Panther Fusion® specimens, respectively (Figure 4).

**TABLE 3.** Culture positivity among PCR-positive specimens subjected to fungal culture.

| Platform | Culture Agreement (95% CI) |
| --- | --- |
| ABI PCR Result | 76.4% (74.4-78.2) |
| BDMAX | 68.1% (65.5-70.6) |
| Fusion | 87.2% (85.2-88.9) |

**FIG 4.**
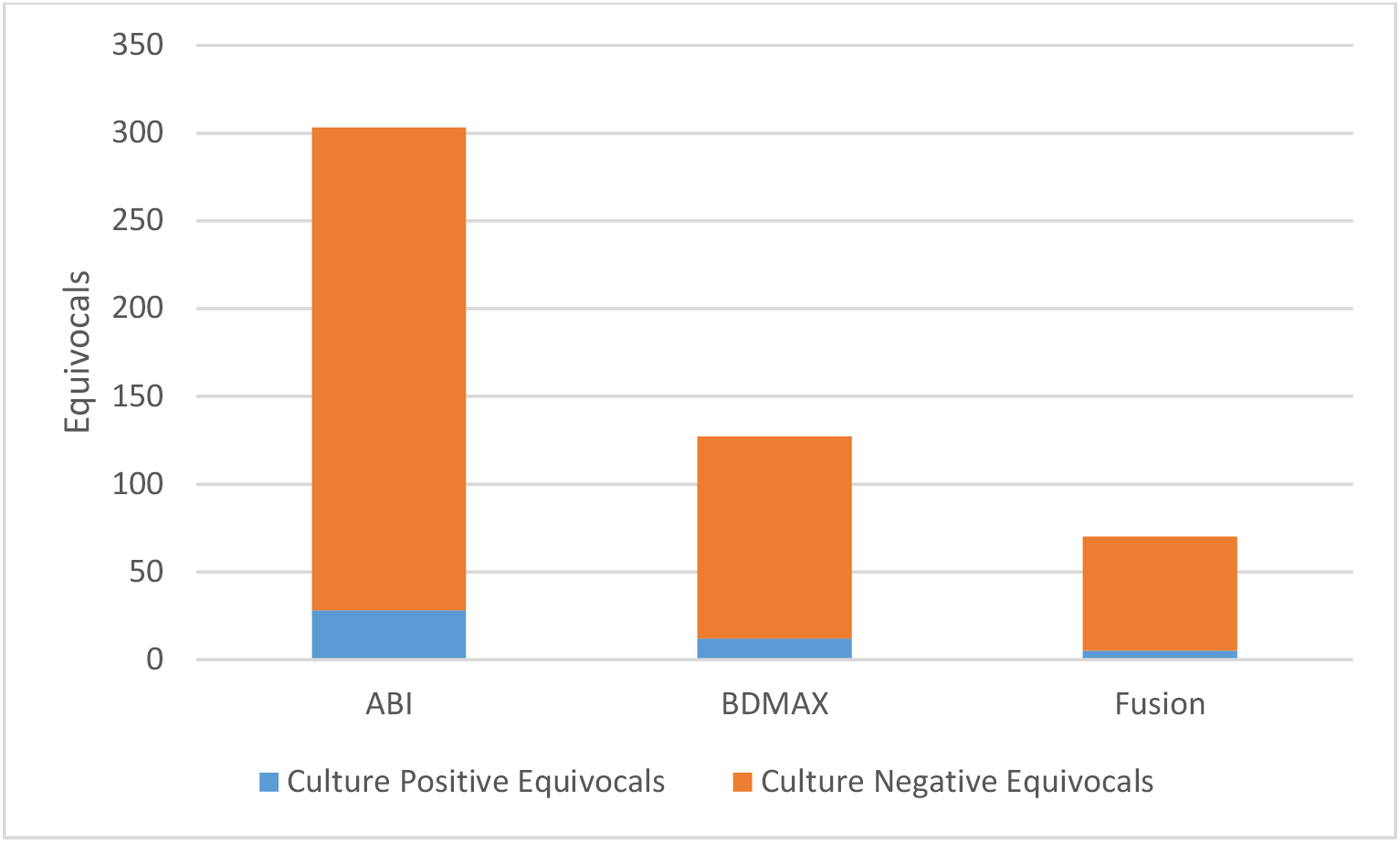
Equivocal culture agreement between ABI, BDMAX, and Fusion platforms.

Because the three platforms were used during different time periods and tested different surveillance populations, retrospective data were not used to estimate comparative clinical sensitivity or specificity.

Implementation of the Panther Fusion® assay increased testing capacity from approximately 88 specimens per work shift on the BD MAX™ platform to as many as 500 specimens per work shift.

## DISCUSSION

*Candida auris* surveillance continues to expand in the United States as healthcare facilities and public health agencies respond to increasing numbers of colonized and infected patients. Rapid identification of colonized individuals is essential for implementation of infection prevention measures, outbreak containment, and regional surveillance efforts. In this study, we developed and validated a laboratory-developed real-time PCR assay for detection of *C. auris* colonization on the Hologic Panther Fusion® Open Access platform and demonstrated its successful implementation within a high-volume regional public health laboratory.

The assay demonstrated strong analytical performance across all validation studies. Agreement with the existing BD MAX™ assay was excellent, with an overall agreement of 97% and a Cohen’s κ value of 0.94. The assay also demonstrated reproducible performance across operators and testing days, detected representatives of all five major *C. auris* clades, and exhibited no detectable cross-reactivity with non-target organisms. Analytical sensitivity was comparable to that reported for previously described molecular assays for *C. auris* surveillance, including the original Wadsworth Center assay and more recent laboratory-developed PCR methods. These findings support the use of the Panther Fusion® platform as an accurate and reliable method for detection of *C. auris* colonization. (7,8,10)

The principal advantage of the Panther Fusion® platform was its substantial increase in testing capacity. As the Midwestern regional laboratory within the CDC Antimicrobial Resistance Laboratory Network, WSLH processes surveillance specimens originating from multiple states and healthcare systems. Increasing specimen volume has challenged laboratories performing *C. auris* screening nationwide, particularly as surveillance recommendations and screening activities have expanded. The Panther Fusion® platform increased testing capacity from approximately 88 specimens per shift on the BD MAX™ system to as many as 500 specimens per shift while maintaining excellent analytical performance. This increase in throughput allows laboratories to accommodate large-scale screening initiatives without proportional increases in staffing or instrument infrastructure.

In addition to increasing throughput, implementation of the assay was associated with reductions in equivocal and indeterminate result frequencies. Although these result categories represented a small proportion of total tests across all platforms, minimizing nondefinitive results is operationally important because such specimens often require repeat testing, additional review, or reflex culture. Reduced frequencies of equivocal and indeterminate results therefore improve workflow efficiency and facilitate more rapid reporting of surveillance results. These benefits are particularly valuable in public health laboratory settings where testing volume can fluctuate substantially during outbreak investigations or regional screening initiatives.

Retrospective analysis demonstrated higher culture recovery among Panther Fusion® PCR-positive specimens than among specimens tested on the ABI and BD MAX™ platforms. However, these findings should be interpreted cautiously. The three platforms were used during different time periods and served different surveillance populations, and culture was not routinely performed on PCR-negative specimens. Consequently, these datasets cannot be used to estimate clinical sensitivity, clinical specificity, positive predictive value, or negative predictive value for the individual platforms. Similarly, differences in positivity rates likely reflect changes in disease prevalence, participating facilities, specimen collection practices, and screened populations over time rather than intrinsic differences in assay performance. For these reasons, the retrospective analysis is best viewed as an assessment of real-world implementation characteristics rather than a direct comparison of diagnostic accuracy.

Our findings complement previous reports describing molecular approaches for *C. auris* surveillance testing. The Wadsworth Center assay established the value of PCR-based screening during large outbreak investigations, and subsequent studies have demonstrated the feasibility of implementing *C. auris* molecular assays across a variety of clinical and public health laboratory settings. The current study extends this work by demonstrating implementation on a highthroughput automated platform within a regional public health reference laboratory responsible for large-scale surveillance testing across multiple jurisdictions. The combination of automated extraction, high throughput, and open-access assay configuration allows laboratories to maintain flexibility while meeting increasing testing demands.

One limitation identified during implementation was that Panther Fusion® specimen lysis buffer did not completely inactivate *C. auris*. Although no laboratory exposures occurred, this observation highlights the importance of maintaining appropriate biosafety practices when handling surveillance specimens. Standard precautions, including proper specimen handling and routine environmental decontamination with agents active against *C. auris*, remain necessary throughout testing workflows.

In conclusion, the Panther Fusion® *C. auris* assay demonstrated excellent analytical performance and successfully supported high-volume surveillance testing within a regional public health laboratory. Implementation substantially increased testing capacity while maintaining accurate and reproducible detection. As demand for *C. auris* screening continues to grow, highthroughput molecular platforms such as Panther Fusion® may play an increasingly important role in supporting regional and national surveillance programs.

## ACKNOWLEDGMENTS

The study was supported by the CDC Epidemiology and Laboratory Capacity (ELC) project I funding (CK24-0002).

